# Comparing the efficacy of cancer therapies between subgroups in basket trials

**DOI:** 10.1101/401620

**Authors:** Adam C. Palmer, Deborah Plana, Peter K. Sorger

## Abstract

An increase in the number of targeted anti-cancer drugs and growing genomic stratification of patients has led to the development of basket clinical trials in which a single drug is tested simultaneously in multiple tumor subtypes under a master protocol. Basket trials typically involve few patients per type, making it difficult to rigorously compare responses across types. We describe the use of permutation testing to analyze tumor volume changes and Progression Free Survival across subtypes in basket trials for neratinib, larotrectinib, pembrolizumab, and imatinib. Permutation testing is a complement to the standard Simon’s two-stage binomial approach and can test for differences among subgroups using empirical null distributions while controlling for multiple hypothesis testing. This approach uncovers examples of therapeutic benefit missed by a binomial test; in the case of the SUMMIT trial, our analysis identifies an overlooked opportunity for use of neratinib in lung cancers carrying ERBB2 Exon 20 mutations.

## INTRODUCTION

In a traditional clinical trial of a cancer therapy, the agent is tested in patients defined by specific inclusion and exclusion criteria that usually involve tissue of origin and disease stage. Growing research and use of molecularly targeted therapies has driven interest in evaluating multiple patient populations with different tumor types, potentially differing in the status of one or more biomarkers (most commonly a genetic alteration). As a result, ‘master protocol’ trial designs have been developed in which several therapeutic hypotheses are tested at the same time via multiple parallel sub-studies (‘baskets’) under a single clinical protocol (and its associated ethical and regulatory reviews). Use of master protocols is intended to expedite the development of drugs by reducing the time and number of patients required to find an efficacious therapy for specific subgroup (Hirakawa et al., 2018; Park et al., 2019; Renfro and Mandrekar, 2018). For example, the NCI-MATCH phase II precision medicine trial (ClinicalTrials.gov number NCT02465060) currently underway is comparing ∼40 treatment arms and multiple genetic biomarkers using a master protocol (Mullard, 2015). Basket trials are particularly helpful when: (i) expanding from an initially successful indication to one or more additional tumor types (ii) searching for a responsive setting in which to perform pivotal trials (iii) studying the predictive value of a biomarker in multiple cancer types (Redig and Jänne, 2015; Tao et al., 2018; Woodcock and LaVange, 2017).

Two recently completed trials demonstrate the potential for basket trials to identify tissue-agonistic biomarkers. When the TRK inhibitor larotrectinib was tested in a diverse set of 12 solid tumors types (NCT02122913, NCT02637687, and NCT02576431) (Drilon et al., 2018) the presence of a TRK fusion gene, irrespective of tumor tissue of origin, was found to identify tumors responsive to larotrectinib. Similarly, in 12 tumor types, mismatch repair (MMR) deficiency was found to be predictive of responsiveness to the PD-1 immune checkpoint inhibitor pembrolizumab (NCT01876511) (Le et al., 2017). In most cases however, both biomarker status and tissue of origin have an influence on drug activity; for example BRAF inhibitors are much less effective in BRAF-mutant colorectal carcinomas than BRAF-mutant melanomas (Korphaisarn and Kopetz, 2016). For any single gene, the type of mutation (i.e.: inhibitory, truncating or activating) can also affect response (Tao et al., 2018). Depending on the way subtypes are defined, a basket trial can be used to assess the impact of one or more of these variables.

The ongoing SUMMIT trial is testing the activity of the ERBB kinase inhibitor neratinib in 21 types of cancer having 42 different mutations in the ERBB2 and ERBB3 receptor tyrosine kinases (*HER2* and *HER3*) (Hyman et al., 2018). Neratinib is an irreversible pan-ERBB (pan-HER) inhibitor approved in 2017 for a relatively narrow indication: patients with early-stage HER2-positive breast cancer who had post-surgical adjuvant therapy using the ERBB2 inhibitor trastuzumab (Singh et al., 2018). Mutation or overexpression of ERBB receptors is implicated in a range of human cancers but ERBB biology is complex, and pre-clinical models provide conflicting data on the potential efficacy of ERBB inhibition in human disease. The multi-center SUMMIT basket trial seeks to resolve this issue by testing neratinib in a wide range of tumor types and genotypes.

In common with a majority of Phase II clinical trials, SUMMIT has no comparator control arm, and SUMMIT instead makes use of a Simon two-stage optimal design to evaluate drug activity based on a dichotomous response metric (Simon, 1989). In this approach, drug response is measured using a radiological assessment of tumor volume according to RECIST criteria (Eisenhauer et al., 2009). Patients whose tumors shrink by ≥30% are scored as responders and others as non-responders; the fraction of responders is the Overall Response Rate (ORR). A binomial test is then used to statistically evaluate the ORR. Using a pre-specified ORR for lack of efficacy (the null hypothesis; typically set at ORR ≤ 10%), the ORR expected under the alternative hypothesis (typically 30%), and the desired rates of Type I and Type II error (≤ 5% and ≤ 20% respectively, corresponding to ≥80% power) the Simon design uses a binomial distribution to calculate the minimum number of patients who must respond in each subgroup for the null hypothesis to be rejected; this calculation is performed separately for each subgroup. If the number of responses in the first stage of a basket is consistent with the null hypothesis, then the treatment is considered futile and corresponding trial arm is terminated. Otherwise the arm expands in a second stage involving additional patients with the goal of testing the alternative hypothesis (e.g. 30% ORR); parameters of the trial design determine the number of patients enrolled in the second stage and the number of responses needed for a therapy to be considered efficacious. The Simon design thereby seeks to detect strong responses in the first phase while minimizing the number of patients subjected to ineffective treatments; it then expands potentially positive subgroups for a larger and more rigorous test in the second phase. In the case of the SUMMIT trial, up to seven patients were initially enrolled per subgroup in Stage 1 and response was evaluated radiologically. Enrollment in each basket was expanded in Stage 2, typically to include 25 patients in total, only if at least one Stage 1 patient exhibited an objective overall response.

Because all basket trials described to date use ORR, in which the assessment of response is dichotomous, the magnitude of tumor volume changes, and changes in the rate of tumor progression, are not considered. The Simon design, as well as Bayesian and frequentist interim analyses developed to determine whether to close enrollment in any subgroups (Cunanan et al., 2017a, 2017b; Drilon et al., 2018; Hyman et al., 2015; LeBlanc et al., 2009; Simon et al., 2016) also assess efficacy *independently* for each subgroup thereby answering the question “*which cancer subtypes surpass a pre-specified threshold for response.”* Here we propose a complementary approach in which tumors are compared across subtypes in a basket trial by using permutation testing to evaluate two related null hypotheses: ‘*no difference in efficacy by tumor type*’ or ‘*no difference in efficacy by class of mutation*’. These hypotheses are plainly relevant to basket trials that may ultimately lead to the approval of therapies for multiple tumor types defined by genetic features. The formulation of hypotheses in this manner has the substantial benefit that all patients enrolled in a trial contribute to the null distribution, and that continuous response variables rather than dichotomous scores can be evaluated (in the current work, duration of Progression-Free Survival (PFS) and magnitude of change in tumor volume). For any specific subgroup, null distributions having an appropriate number of patients for each subgroup are generated by subsampling the all-patient distribution. When response rates are low, as in SUMMIT, the ‘*no difference*’ null hypothesis is similar to a null hypothesis of ‘*low or no activity*’ and can be used to test whether any group has significantly superior responses; when response rates are high, as with larotrectinib, the ‘*no difference*’ hypothesis tests for both inferior and superior responses. In the case of SUMMIT, lung cancers fail Simon criteria but significantly exceed the no-difference null with respect to volume changes and PFS. In contrast, breast cancers in SUMMIT exhibit a high ORR, but are no different from average with respect to PFS. These data suggest an alternative approach for interpreting basket trials with the potential to better discover therapeutic opportunities for subsequent testing in Phase III trials.

## RESULTS

### Analysis of SUMMIT trial reveals overlooked therapeutic opportunity for neratinib in lung cancers carrying ERBB2 Exon 20 mutations

Results for the first 141 patients in the SUMMIT basket trial were recently reported (Hyman et al., 2018). Multiple genetic markers were assessed, including 31 unique HER2 and 11 unique HER3 mutations. Clinical response was measured by radiological assessment of tumor volume changes and by progression free survival (PFS), the time from enrollment until death or radiological evidence of tumor progression. FDA guidance recommends the use of ORR as measured by RECIST criteria (Eisenhauer et al., 2009) in master protocol trials (Research, 2019) largely because ORR is an accepted surrogate endpoint for accelerated drug approval (Pazdur, 2008). Although the SUMMIT trial uses ORR, the authors report changes in tumor volume as a continuous variable.

In common with previous basket trials (Cunanan et al., 2017b) SUMMIT (Hyman et al., 2018) recorded PFS data but it was not analyzed formally or compared to ORR; this reflects the perceived challenge of evaluating 21 tumor types using data from only 141 patients. An additional concern is that PFS duration may not be comparable for cancers having different rates of progression. However, it is also controversial whether tumor volume changes are predictive of overall survival (OS; the ‘gold standard’) (Buyse et al., 2000; El-Maraghi and Eisenhauer, 2008; Fleming and DeMets, 1996; Kaiser, 2013). For example, in a retrospective analysis of non-small cell lung cancer, PFS was correlated with OS (Blumenthal et al., 2015) but ORR was not. The use of PFS in breast cancer trials is also supported by a variety of other data (Adunlin et al., 2015). Thus, although it is standard practice to rely on ORR not PFS in basket trials, we hypothesized that use of both of types of information might provide new therapeutic insights (see Discussion). There is no established method for thresholding PFS data into dichotomous responder and non-responder classes. Thus, it is not possible to use a binomial test. We instead used permutation testing by using repeated Monte Carlo resampling of the distribution of continuous volume changes and PFS from all patients to construct null distributions for each subgroup. The null hypothesis in this case is that there is no difference in volume change or PFS for the subgroup (defined by tumor type or genotype) relative to all patients following exposure to neratinib.

When neratinib-treated patients in SUMMIT were classified by tissue of origin (Figure 1A) and compared to an appropriately resampled ‘*no difference*’ null distribution, breast cancers exhibited significantly greater volume reduction than any other tumor type (p<10^-6^; a 45% difference in average volume change from all non-breast tumors). This agrees with the conclusion by Hyman *et al* that breast cancers are the most neratinib-responsive of all tumor types tested based on ORR (Hyman et al., 2018). Because breast cancers dominate volume-change data we constructed a second set of null distributions for volume changes that included only non-breast tumors (hereafter NB; see Methods).

**Figure 1.**
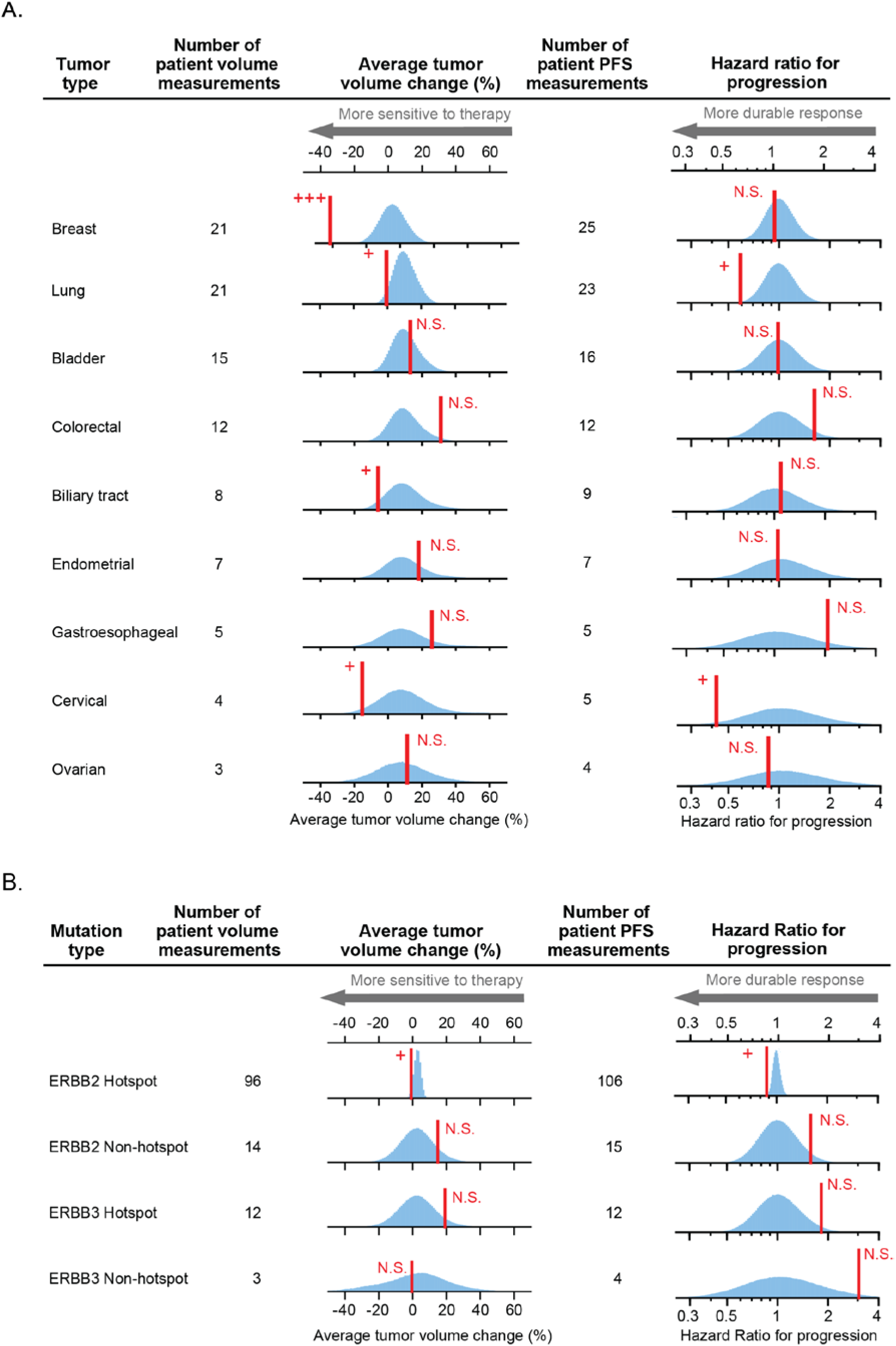
Analysis of neratinib response by tumor tissue of origin and mutation. Red line: observed response. Blue histogram: responses simulated according to the null hypotheses of *no difference* in response between tumors types (A) or mutation types (B). As explained in the main text, breast tumor volume changes are compared with null distributions drawn by Monte Carlo resampling from all tumors; for this reason, the null distribution for breast tumor volume changes has a different mean. For all other tumor volume changes, the null distributions are drawn from all non-breast tumors due to breast tumors being a strong outlier (p < 10^-6^; see Methods). ‘Hazard ratio for progression’ null distributions are drawn from all tumors. Observed responses that significantly exceed the null hypothesis, according to Benjamini-Hochberg procedure for multiple hypothesis testing, are indicated with +; N.S. denotes not significant; +++ denotes p < 10^-6^ (Supplementary Tables S1, S2, S3, S4).

When NB distributions were resampled and compared to tumor-specific volume change data, lung, cervical, and biliary cancers were found to significantly exceed the ‘*no difference by type*’ null hypothesis (*P*=0.04, 0.04 and 0.06; significant according to Benjamini-Hochberg procedure; Supplementary Table S1A). Whereas cervical and biliary cancers passed the criteria for the first stage of a Simon two-stage design, lung cancer failed at the second stage (Table 1). Thus, quantitative criteria judge as positive a volume change in lung cancers that was found to be negative by the binary criteria used in a two-stage design. This discordance arises because half of lung cancers shrank on therapy but only one shrank enough to be classified as a response by RECIST. The permutation test and Simon criteria therefore provide different insights into the drug responsiveness of this small patient population.

**Table 1.** Conclusions from analysis of neratinib in ERBB-mutant tumors in context of trial status.

| Tumor type | Number of patients | Status in Simon Optimal 2-stage design |  | Responses significantly different from other tumors<sup>1</sup>? |  |
| --- | --- | --- | --- | --- | --- |
|  |  | Stage 1 | Stage 2 | Volume | PFS |
| Ovarian | 4 | Ongoing |  | - | - |
| Gastroesophageal | 5 | Ongoing |  | - | - |
| Colorectal | 12 | Failed |  | - | - |
| Bladder | 16 | Failed |  | - | - |
| Endometrial | 7 | Failed |  | - | - |
| Biliary | 9 | Passed | Ongoing | Superior | - |
| Cervical | 5 | Passed | Ongoing | Superior | Superior |
| Lung | 26 | Passed | Failed | Superior | Superior |
| Breast | 25 | Passed | Passed | Superior | - |

<sup>1</sup> Dash denotes no significant difference by Benjamini-Hochberg procedure.

### Analysis of Progression Free Survival

Comparison of response duration among different types of tumors is potentially complicated by differences in tumor kinetics. While slow growth should not in and of itself equate to ‘sensitivity’ to therapy, durability of response is clinically important, is commonly an endpoint in cancer trials, and may provide orthogonal data to complement measurement of volume change. We therefore applied permutation testing to PFS. The null distribution was drawn from all tumor types (n=141) because no tumor type was so responsive as to dominate the distribution (Methods). Significantly smaller hazard ratios, which are indicative of longer PFS, were identified by a *no difference* test in cervical cancers (*P*=0.03; median PFS 20 months) and lung cancers (*P*=0.003; median PFS 5.4 months) but - strikingly - not in breast cancers (*P*=0.36; median PFS 3.5 months, Supplementary Table S1B). Only five neratinib-treated cervical cancers are present in the SUMMIT dataset, and the empirical null distributions was consequently broad (Figure 1A). Nonetheless, the observed responses were sufficiently strong and durable to achieve statistical significance (cervical tumors also met the criteria to begin Stage 2 and so additional patients are currently accruing; Table 1). Whereas lung cancers exceed *no difference* tests for both volume changes and hazard ratios, breast cancers differ from the overall population by volume change alone. Lung cancers therefore appear to represent a therapeutic opportunity for neratinib missed by dichotomous assessment of response.

Our approach identifies differences in PFS that are statistically significant, but interpreting whether this is clinically meaningful requires attention to absolute duration in context of the kinetics of that specific tumor type. In this case, as noted by Hyman, a therapeutic response exceeding 12 months in non-small cell lung cancer is clinically meaningful (Hyman et al., 2018). Moreover, in the case of neratinib-treated lung and cervical cancers, significant differences from the null distribution were observed for both volume change and PFS data, increasing confidence in the conclusions (see also Discussion).

### Analysis of biomarkers

Differences in neratinib sensitivity have been observed in cell lines with different mutations in ERBB receptors (Nagano et al., 2018) but the impact of such differences has not been reported for patients. Basket trials provide the opportunity to investigate this issue. SUMMIT enrolled patients on the basis of qualifying mutations in ERBB2 or ERBB3, which were classified as ‘hotspot’ if they occurred in recurrently mutated regions of either gene, or ‘non-hotspot’ if they lay in rarely mutated regions (Hyman et al., 2018). However, conducting independent two-stage trials in multiple tissue types results in evaluating responsiveness and cohort expansion on the basis of tumor tissue of origin alone without considering the influence of genotype. We therefore applied permutation testing to ERBB genotypes and neratinib responses. We found that tumors with ERBB2 hotspot mutations exceeded the *no-difference* null model as judged by changes in tumor volume or PFS (Figure 1B) (*P*=0.0005 for PFS and *P*=0.03 for volume changes), which agrees with Hyman’s conclusion that ERBB2 hotspot tumors are responsive to therapy. When ERBB2 hotspot mutations were further divided into functional classes (e.g. S310; Exon 20 insertions; V777; L755; and “other hotspot mutations”), Exon 20 insertions significantly exceeded the *no difference* null for PFS (*P*=0.01), which could be attributed almost exclusively to lung tumors (Hyman et al., 2018) (6 lung tumors were among the 7 most durable responses observed for all cancer types having Exon 20 insertions). No other significant signals were detected among subgroups when scoring for mutation class.

### Permutation testing provides statistical support for the use of imatinib only in select cancer types

As a second application of our approach we examined the phase II open label *Imatinib Target Exploration Consortium Study B2225* which tested Imatinib in 186 patients having 40 different malignancies (Heinrich et al., 2008). Objective responses were observed in six types of malignancy, of which five were described as “notable” but not subjected to formal statistical analysis. By testing against a *no difference* null we found that three malignancies had a significantly higher ORR to imatinib than all other tumors tested (dermatofibrosarcoma protuberans, myeloproliferative disorders, hypereosinophilic syndrome; Supplementary Table S4A). These malignancies were represented by 7 to 14 patients each, out of 186, confirming that statistically significant drug activity can be detected in small subgroups within a basket trial. Imatinib was approved for use in dermatofibrosarcoma protuberans by the FDA in 2006, and, following a phase 2 study published in 2010 (NCT00122473) it was incorporated into the National Comprehensive Cancer Network’s treatment guidelines for this malignancy (Navarrete-Dechent et al., 2019). The use of imatinib in hypereosinophilic syndrome is supported by case studies (Gleich et al., 2002; Pardanani and Tefferi, 2004) and our analysis demonstrates additional support in a Phase II basket trial (Heinrich et al., 2008).

### Permutation testing provides statistical support for tumor-agnostic use of larotrectinib and pembrolizumab in biomarker positive populations

Basket trials of the immune checkpoint inhibitor pembrolizumab (Le et al., 2017) and kinase inhibitor larotrectinib (Drilon et al., 2018; Lassen et al., 2018) contrast with trials of neratinib and imatinib because response rates are much higher: both drugs were found to be effective in multiple types of tumors positive for a specific genetic biomarker. In a basket trial of 86 patients and 12 tumor types, tumors with mismatch repair (MMR)-deficiency were found to be highly responsive to PD-1 blockade by pembrolizumab regardless of tissue of origin (Le et al., 2017). Similarly, high rates of larotrectinib response were observed among 122 patients having 15 different types of tumors expressing TRK fusion proteins (Drilon et al., 2018; Lassen et al., 2018). When data from each of these trials were compared to a *no difference* null (testing in for both superiority and inferiority), no significant differences were observed for any tumor type represented by three or more patients (this corresponded to eight tumor types for larotrectinib and seven types for pembrolizumab). The sole exception was infantile fibrosarcomas, which were more responsive to larotrectinib than other TRK-fusion tumors (Figures 2A, 2B, Supplementary Tables S4B, S4C). Thus, a comparison among tumor types provides additional statistical support for the conclusion that larotrectinib and pembroluzimab exhibit tumor-agnostic activity in tumors having specific genetic features.

**Figure 2.**
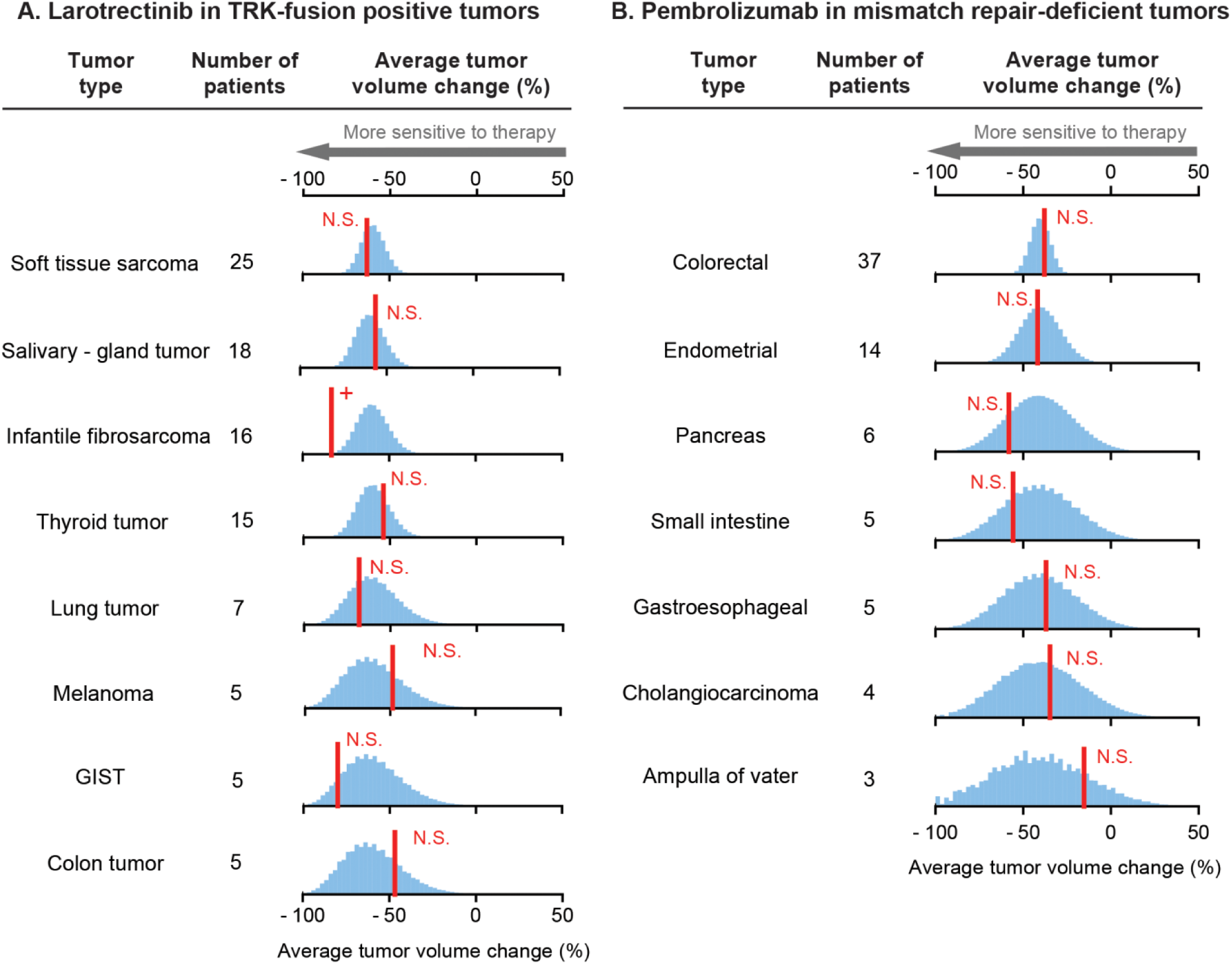
Analysis of larotrectinib and pembrolizumab responses by tumor tissue of origin. Red line: observed average response. Blue histogram: responses simulated according to the null hypothesis of *no difference* in response between tumors types. Observed responses that significantly exceed the null hypothesis, according to Benjamini-Hochberg procedure for multiple hypothesis testing, are indicated with +; N.S. denotes not significant (Supplementary Table S4).

### Comparison of Type 1 and Type 2 errors of permutation tests and binomial tests in basket trials

When some but not all tumor subtypes respond to therapy, the responsive subtypes may potentially be identified by either of permutation tests (which evaluate a ‘*no difference by tumor type*’ null using continuous measures of responses) and binominal tests as those in the Simon two-stage design (which evaluate a ‘*low efficacy*’ null independently for each tumor type using dichotomous measures of responses). To compare rates of type I error (a false positive corresponding to misclassification of a non-responsive tumor type as responsive) and type II error (a false negative, corresponding to misclassification of a responsive tumor type as non-responsive) between these approaches, we simulated basket trials in which a varying proportion of tumor subtypes responded to therapy (Supplementary Methods S6). As expected, by permutation testing on continuous volume change data, the rate of type I error declined as the treatment effect increased (i.e. the decrease in tumor volume was greater). In small cohorts typical of the first stage of a two-stage trial (N=7 patients per tumor type), permutation tests had substantially smaller type I error than binomial tests. This is potentially important because a key aim of two stage trial designs is to minimize patients exposed to futile treatments (Figure 3A). In larger cohorts typical of Stage Two (N=25 patients per tumor type), permutation tests had greater power for all effect sizes than a binomial test, as well as smaller type I error for treatment effects stronger than 20% difference in tumor volume (Figure 3B). Thus permutation testing appears to be the more powerful approach, in agreement with recent theoretical analysis (Arfé et al., 2019). Historically, an important advantage of binomial tests was that they could be computed rapidly and exactly with simple algorithms and slow computers. Permutation testing with re-sampling (necessary when N is too large for an exact enumeration) is more computationally intensive; this was an issue in 1980s when basket trials were first proposed, but it is no longer relevant.

**Figure 3.**
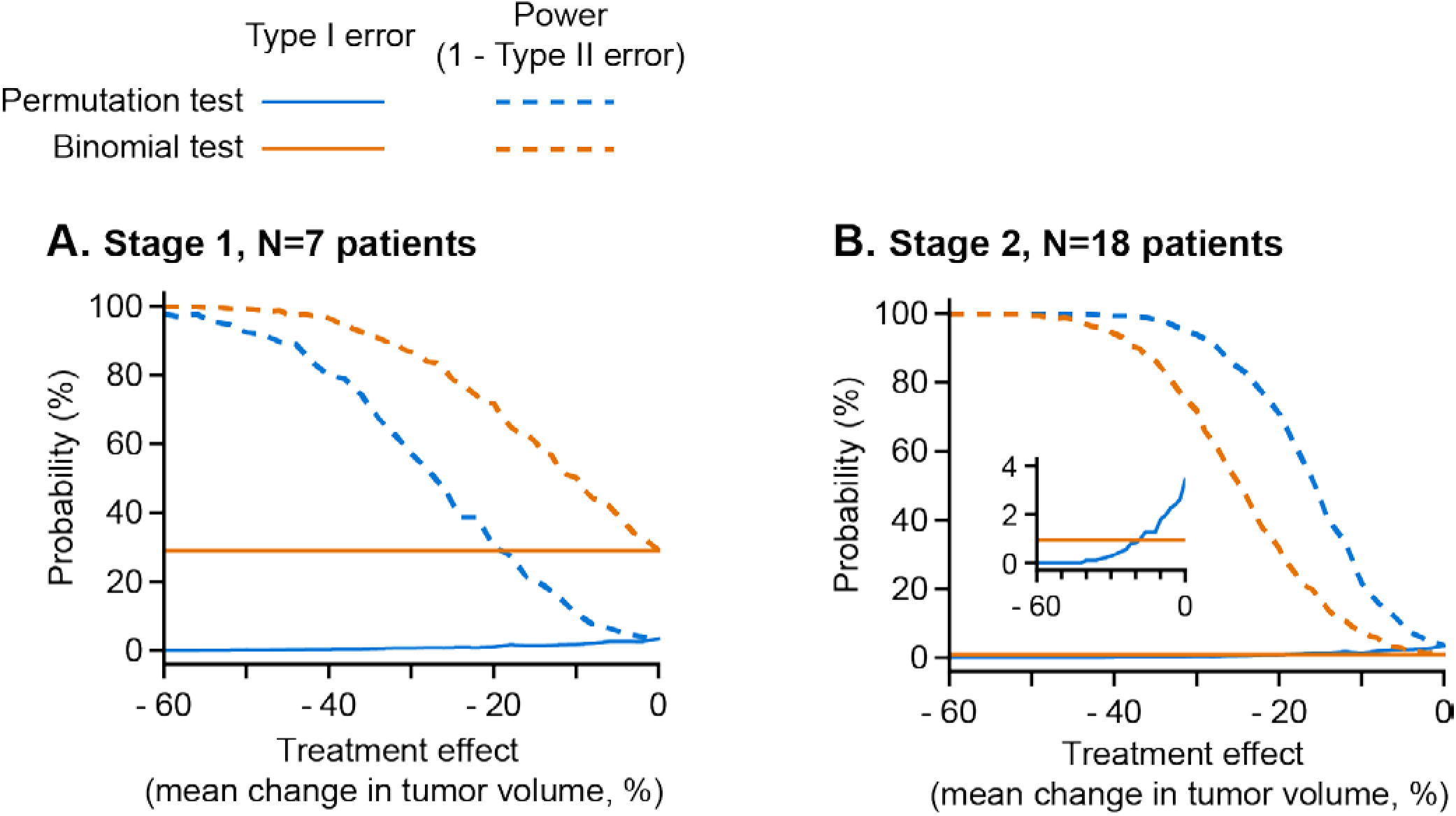
Comparison of Type 1 and Type 2 errors of permutation tests and binomial tests in basket trials. Basket trials were simulated in which three of ten tumor types respond to therapy, and sensitivity and specificity (Type 1 and Type 2 errors) were compared between: (blue) permutation tests, comparing all tumor types to find those significantly more responsive than average, and (orange) binomial tests of objective response rate, such as are used in Two-Stage trial designs (see Supplementary Method). Simulations were repeated across a range of treatment effect sizes (difference in mean volume change between responsive and non-responsive tumors) for 7 patients per tumor type (**A**, typical of the first stage of a two-stage trial), and 18 patients per tumor type (**B**, typical of the second stage). **B**, inset: zoom on the type 1 error rate (<4%).

## DISCUSSION

A primary motivation for performing a basket trial is to determine which of several tumor types or genotypes are sufficiently responsive to an investigational therapy to warrant further study in a Phase III pivotal trial. Because Phase II trials rarely involve a no-treatment control population, contemporary designs for basket trials use a pre-specified cutoff to evaluate whether or not a drug is effective. Currently this involves a dichotomous assessment of tumor volume changes to determine if the overall response rate exceeds a threshold set by a binomial test. In this paper we demonstrate an alternative approach involving a permutation test in which both continuous volume changes and survival data (PFS) are formally compared against empirical null distributions constructed using data from all patients in the trial. Responses in subgroups are then compared to the null distribution to test the hypothesis of *no difference in efficacy by subtype* (most commonly tumor tissue of origin or mutation class or genotype) as a means to identify subtypes that are most responsive.

Constructing subtype-specific null distributions involves repeated Monte Carlo resampling of an all-patient distribution, drawing the same number of samples as the number of patients in the subtype. The resulting null distributions appropriately anticipate the greater variability observed in small cohorts, thereby adjusting the threshold for identifying a statistically significant increase or decreases in response based on a pre-specified type 1 error rate. For example, the SUMMIT trial reported PFS data for five cervical cancer patients. In this case, the null distribution was calculated by repeatedly sampling five response durations from the set of duration data for all patients, generating a relatively wide subtype-specific null distribution. Despite this, the observed hazard ratio in cervical cancers was significantly smaller than the *no difference* null distribution (*P*=0.03) implying an above-average response. Conclusions drawn from testing for *no difference* in continuous volume change can differ from binomial testing based on ORR. For example, lung cancers exposed to neratinib exceed the *no difference* null with respect to both volume changes (*P*=0.04; sampling from all non-breast tumors) and PFS (*P*=0.003, sampling from all tumors) even though lung cancers failed the second stage of a Simon design. In contrast, breast cancers exhibited highly significant changes in tumor volume by both Simon and *no difference* criteria, but failed the *no difference* test with respect to PFS. We therefore propose that neratinib be studied further in ERBB-mutant lung tumors and that early evidence be sought in expansion cohorts of whether neratinib is providing a clinically meaningful survival benefit in breast cancer patients.

Basket trials of larotrectinib in TRK fusion-positive cancers and pembrolizumab in MMR-deficient cancers are characterized by high response rates (Drilon et al., 2018; Lassen et al., 2018; Le et al., 2017). By permutation testing, no subgroup was identified in either trial that was significantly less drug-responsive than the average of all tumors. Thus, a formal *no difference* test supports the recent tumor-agnostic FDA approvals of larotrectinib and pembrolizumab for cancers with specific genetic features.

### Comparison of subgroups in basket trial

It is not conventional to directly compare responses across arms of a basket trial and FDA guidance discourages this, probably because of the dangers of multiple hypothesis testing (Pazdur, 2008; Research, 2019). The specific concern is that, in trials with a large number of arms, testing all arms against each other involves a potentially uncontrolled multi-hypothesis test. However, in the procedure described here, all null distributions are sampled from the same all-patient distribution and the Benjamini-Hochberg procedure is used to appropriately correct the significance threshold used to test hypotheses. In some cases one tumor subtype can dominate responses for the entire trial, obscuring smaller but potentially significant differences in other subtypes. In SUMMIT this was observed for volume changes in neratinib-treated breast cancers (*P*<10^-6^ relative to the *no difference* null). To enable detection of next-most different volume responses, we removed breast cancers from the all-patient distribution. We performed this procedure only for a single outlier subgroup because repeated adjustment of the null distribution heightens the risk of false discovery from multi-hypothesis testing (Bishop and Thompson, 2016) (see Methods).

A second concern arises when comparing subgroups directly: because tumors respond differently to therapy, the magnitude of volume changes and the frequency of confounding factors such pseudo-progression (an increase in the size of a primary tumor for reasons other than disease progression, such as immune infiltration, followed by tumor regression) (Ma et al., 2019) also differ. However, in scoring ORR in the Simon design, a very similar issue arises: the same threshold for volume change is used to establish a meaningful response in all subgroups. Thus the need for a common interpretation of subgroups is *not* specific to our Monte Carlo resampling methodology and any other method (Bayesian, frequentist) of assessing drug efficacy (Berry, 2015). It is also noteworthy that the 30% reduction in volume conventionally used to threshold ORR subdivides a unimodal distribution of tumor volume changes. It is justifiable based on the complexities of tumor volume assessment (Sharma et al., 2012) but is nonetheless arbitrary.

A third concern involves our use of PFS data to compare subgroups; this arises because different cancers naturally progress at different rates (Friberg and Mattson, 1997). Empirical data demonstrate, however, that rates of progression for solid tumors in the SUMMIT trial are similar to each other: tumors that did not shrink on therapy progressed rapidly irrespective of tumor type, (85% of non-shrinking tumors progressed in ≤ 3 months). Moreover, it is well established that overall survival, the gold standard for measuring response to anti-cancer drugs, correlates more strongly with duration of PFS than with tumor volume changes (Fleming and DeMets, 1996; Kaiser, 2013; Seymour et al., 2010; Zabor et al., 2016). Significant reductions in tumor volume do not necessarily predict durable PFS, and durable PFS can be achieved with modest changes in tumor volume. Radiological measurement of tumor volume is also complicated by various technical and biological factors. Thus, past experience and theoretical considerations suggest that PFS and tumor volume can both provide valuable data in a permutation testing framework.

Historically, a final limitation in the use of PFS data in basket trials is that there exists no agreed upon threshold in duration that can define a meaningful (or ‘objective’) response; in contrast tumor volume changes are usually thresholded at a common value for determination of ORR. In the absence of a PFS threshold and a dichotomous score a binomial test cannot be used. However, permutation testing using an all-patient null distribution overcomes this issue.

In conclusion, we describe a simple permutation test for small patient populations that makes it possible to obtain appropriately scaled null distributions and derive empirical *P* values for drug response as measured by both volume change and PFS. The methodology is expected to be of value in basket trials and other Phase II studies that lack control arms and involve multiple patient subgroups generally thought to be too small for formal comparison (Hyman et al., 2018). Moreover, among all tests which control the Type I error rate at a fixed α level, the permutation test has been proven mathematically to be the testing procedure that maximizes finite-sample power for a late stage study conditional on early-stage data (Arfé et al., 2019). Thus, use of permutation testing in basket trials is expected to be of greater predictive value for subsequent Phase III studies. Nonetheless, all comparisons of trial subgroups must be interpreted in the context of known differences in tumor growth rates that may affect tumor volume changes and duration of PFS.

The continuing growth of genomic-driven oncology will increasingly enable refined subdivision of patient populations whether in a basket trial or by stratifying patients in conventional Phase II and Phase III studies (Hyman et al., 2018). The promise of such subdivision is better precision in oncology, but the risk is smaller subsamples and reduced statistical significance; thus new approaches are required. Our reformulation of null hypotheses, generation of null distributions by permutation, and derivation of empiric P values for comparing responses across subgroups in basket trials has the potential to better identify therapeutic opportunities for targeted drugs. The approach is grounded less in novel statistical theory (permutation tests are well established) but rather in accumulating empirical evidence from completed basket trials. This study does not in our view necessitate changes in trial designs, rather, we suggest that an appropriately conservative approach is to continue the use of established methods such as the Simon or Bayesian Simon designs for enrollment decisions and evaluations of absolute efficacy, and to apply permutation testing for subsequent analysis of differences in efficacy among subgroups.

## ACKNOWLEDGEMENTS

We thank L. Trippa and B. Alexander for helpful discussions and comments on this manuscript. This work was supported by NIH grants P50-GM107618 and U54-CA225088 (to PKS). D.P. is supported by NIGMS grant T32-GM007753.

## AUTHOR CONTRIBUTIONS

Analysis, A.C.P. and D.P.; writing, A.C.P., D.P., and P.K.S.

## METHODS

To test the null hypothesis that patient subgroups are equally responsive to a therapy, outcome data as reported in a basket trial (comprising either change in tumor volume, or duration of PFS) were pooled for all patients who received the drug, regardless of tumor type. We derive a null distribution for each subgroup by permutation of responses among tumor subgroups.

Exact permutation tests compute all possible combinations of categorical variables, but this is computationally intractable for continuous variables (e.g. there are 10^23^ ways to choose 25 samples from 100 patients). We therefore used Monte Carlo permutation tests, in which a large but non-exhaustive set of permutations is randomly generated. Monte Carlo permutation yields type 1 error rates (false positive rate) equal to those of an exact permutation test for probabilities P >> 1/N where N is the number of random permutations; we used N=10^7^ and therefore can accurately report P values as small as 0.0001 (10^6^ simulations were performed for the neratinib PFS analysis due to the computational time required to calculate hazard ratio, and since neratinib PFS analyses produced no P values smaller than 10^−4^, sufficient precision was provided by 10^6^ simulations). Monte Carlo permutation of trial outcomes involves randomly drawing from a pool of all patient responses, with the number of samples drawn equal to the number of patients found in the cohort being tested (e.g. 26 patients for lung and 5 patients for cervical cancer). A response metric (volume change or PFS) for the sampled set is then calculated and the procedure repeated N=10^7^ times to compose a reliable null distribution for the cohort. For the analysis of changes in tumor volume, the response metric was the average volume change for a cohort; for the analysis of PFS, the response metric was the hazard ratio (computed using the Cox proportional hazards model) of the Kaplan-Meier survival function for a subset of patients as compared to the survival function for all patients. An empiric P value was then determined by the location of the observed response metric (which was the test statistic) on that null distribution. In common with an exact permutation test, the rate of type I error is the significance level. The Benjamini-Hochberg procedure (Benjamini and Hochberg, 1995) was used to control the False Discovery Rate (FDR) associated with multiple hypothesis testing (multiple hypothesis correction is generally absent from analyses of basket trials). Consistent with practice in genomics, we used an FDR of 25%, which we observed by simulations to yield a type I error rate ≈ 3% (see results); this is smaller than the 10% rate of type 1 error commonly chosen for Simon two-stage designs.

In the case of the SUMMIT trial permutation testing was separately applied to reported tumor volume changes and to durations of PFS; in the case of the larotrectinib and pembrolizumab trials (Drilon et al., 2018; Lassen et al., 2018; Le et al., 2017) it was applied only to tumor volume changes (PFS outcomes by tumor type are not available). For imatinib, permutation tests were applied to objective response rates (Heinrich et al., 2008). For the SUMMIT trial, volume (but not PFS) changes in breast tumors were far stronger than for any other tumor type: none of 10^7^ simulations of the null hypothesis matched the observed average tumor volume change of breast tumors (we report this as P < 10^-6^). The magnitude of difference between breast tumors and all tumors (45% difference in average volume change) is so large that the inclusion of breast tumors in the null distribution makes it impossible to detect any difference among other tumor types. Because breast tumors represent an outlier with regard to volume changes in response to neratinib treatment, we considered it inappropriate to include breast tumor volume changes in the between-tumor comparison of all other tumor types. We therefore constructed a “no breast tumor” (NB) null distribution using volume data for all non-breast cancers (n=116). This reformulation of the null distribution was applied only for this case of a *P*<10^-6^ outlier, and we advocate for a similarly stringent approach to any future application that may remove subtypes from the null distribution. We did not encounter any other tumor subtype in any basket trial for which reformulation of the null distribution was appropriate

Responses in any one tumor type could not be meaningfully inferior to the poor response across all patients to neratinib (median volume change ≈ 0%; median PFS ≈ 2 months; objective response rate 12%). We therefore tested only for superiority of each tumor type or mutation class relative to all types; the same was true of imatinib (objective response rate 13% over all patients), and basket trials in general use one-sided tests for efficacy. In the cases of larotrectinib and pembrolizumab, overall response rates were high, and we tested for both superiority and inferiority relative to the average of all tumors in those trials.

Finally, basket trials were simulated in which only some tumor types respond to therapy, in order to compare type I and type II error rates between permutation tests (comparing efficacy across tumor types) and binomial tests (evaluating objective response rate in individual tumor types, according to a Simon Two-Stage trial design). A ‘non-responsive’ distribution of tumor volume changes was empirically defined based on the observed volume changes in non-responsive tumor types in the SUMMIT trial: volume changes were drawn from a normal distribution with mean response μ = +20%, and standard deviation σ = ±30%; these parameters resulted in fewer than 5% of tumors exhibiting volume change ≤ -30%, defined as ‘objective response’ for these simulations. Basket trials were simulated in which ten tumor types were studied, of which seven types were ‘non-responsive’ (μ = 20%, σ = ±30%), and three types were ‘responsive’ (μ = α + 20%, σ = ±30%; where α is the ‘treatment effect’, the average difference in volume change compared to non-responsive tumors). 1000 basket trials were simulated for each value of ‘treatment effect’ between -60% and 0%, first with 7 patients per tumor type, and next with 18 patients per tumor type, matching the intended number of patients in Stages One and Two of the two-stage design of the SUMMIT trial. Each simulated trial’s results were analyzed by both permutation testing, and by the binomial test used in the Two-Stage design (pass requires ≥ 1 objective response at Stage One, and ≥ 4 objective responses at Stage Two). Type 1 error rates were calculated as the fraction of truly non-responsive tumor types that were misclassified as responsive, and Type 2 error rates were calculated as the fraction of truly responsive tumor types that were misclassified as non-responsive.

## SUPPLEMENTARY INFORMATION

**Supplementary Table S1.** Benjamini-Hochberg critical values for analysis of neratinib tumor volume responses (A) and hazard ratios (B) by tumor tissue of origin. Related to Figure 1.

| Tumor Type | Test for larger benefit in tumor volume |  | Conclusion based on 25% False Discovery Rate |
| --- | --- | --- | --- |
|  | p-value | Benjamini-Hochberg critical value for 1-sided test |  |
| **Cervical** | **0.039** | **0.031** | **Larger Benefit** |
| **Lung** | **0.040** | **0.063** | **Larger Benefit** |
| **Biliary Tract** | **0.059** | **0.094** | **Larger Benefit** |
| Ovarian | 0.569 | 0.125 | N.S. |
| Bladder | 0.659 | 0.156 | N.S. |
| Endometrial | 0.768 | 0.188 | N.S. |
| Gastroesophageal | 0.872 | 0.219 | N.S. |
| Colorectal | 0.977 | 0.250 | N.S. |

B.
| Tumor Type | Test for larger benefit in hazard ratio for progression |  | Conclusion based on 25% False Discovery Rate |
| --- | --- | --- | --- |
|  | p-value | Benjamini-Hochberg critical value for 1-sided test |  |
| **Lung** | **0.003** | **0.028** | **Larger Benefit** |
| **Cervical** | **0.027** | **0.056** | **Larger Benefit** |
| Ovarian | 0.347 | 0.083 | N.S. |
| Breast | 0.363 | 0.111 | N.S. |
| Endometrial | 0.454 | 0.139 | N.S. |
| Bladder | 0.467 | 0.167 | N.S. |
| Biliary tract | 0.579 | 0.194 | N.S. |
| Gastroesophageal | 0.912 | 0.222 | N.S. |
| Colorectal | 0.938 | 0.250 | N.S. |

**Supplementary Table S2.**
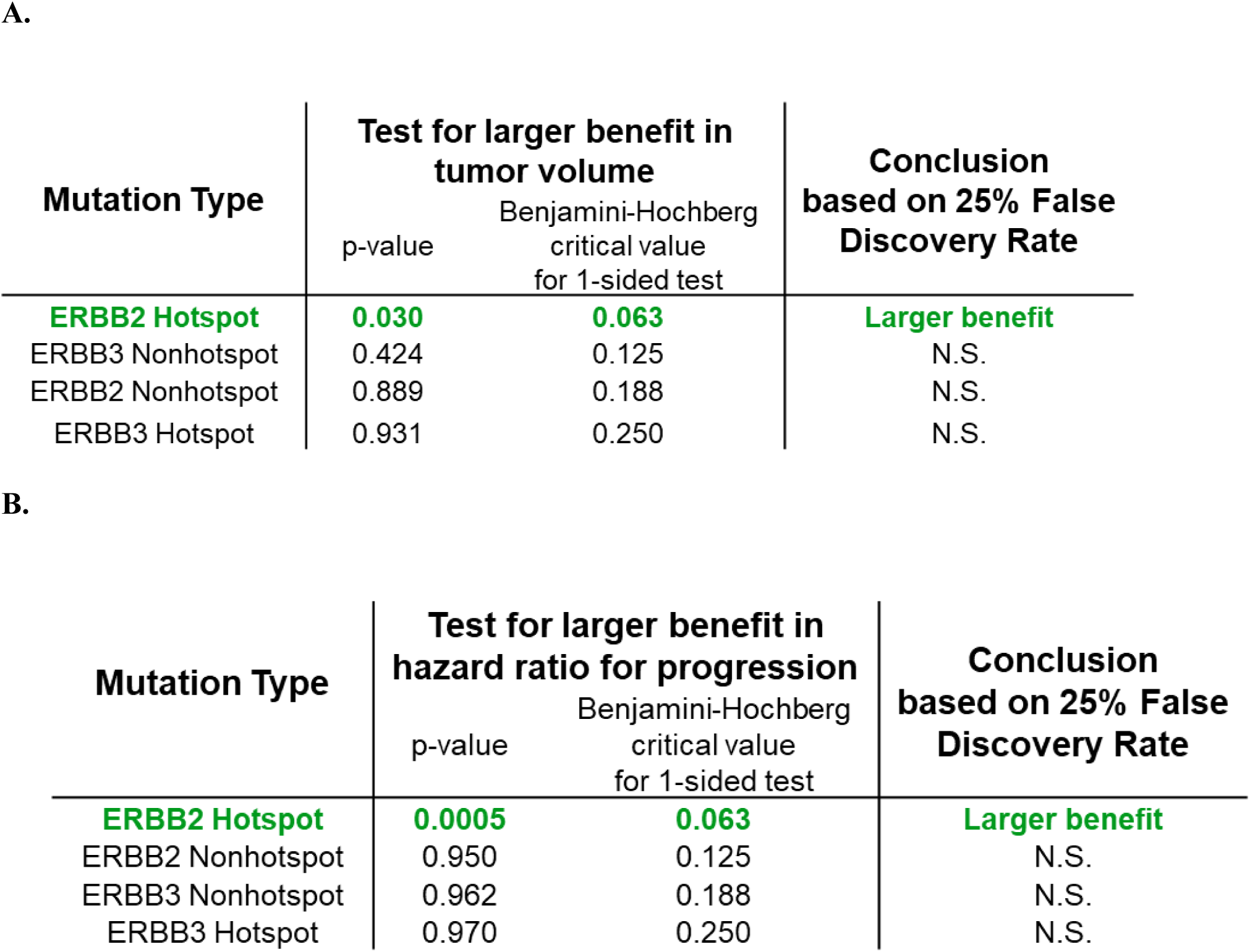
Benjamini-Hochberg critical values for analysis of neratinib tumor volume responses (A) and hazard ratios for progression (B) by general mutation type. Related to Figure 2.

| Mutation Type | Test for larger benefit in tumor volume |  | Conclusion based on 25% False Discovery Rate |
| --- | --- | --- | --- |
|  | p-value | Benjamini-Hochberg critical value for 1-sided test |  |
| **ERBB2 Hotspot** | **0.030** | **0.063** | **Larger benefit** |
| ERBB3 Nonhotspot | 0.424 | 0.125 | N.S. |
| ERBB2 Nonhotspot | 0.889 | 0.188 | N.S. |
| ERBB3 Hotspot | 0.931 | 0.250 | N.S. |

B.
| Mutation Type | Test for larger benefit in hazard ratio for progression |  | Conclusion based on 25% False Discovery Rate |
| --- | --- | --- | --- |
|  | p-value | Benjamini-Hochberg critical value for 1-sided test |  |
| **ERBB2 Hotspot** | **0.0005** | **0.063** | **Larger benefit** |
| ERBB2 Nonhotspot | 0.950 | 0.125 | N.S. |
| ERBB3 Nonhotspot | 0.962 | 0.188 | N.S. |
| ERBB3 Hotspot | 0.970 | 0.250 | N.S. |

**Supplementary Table S3.** Benjamini-Hochberg critical values for analysis of neratinib tumor volume responses (A) and hazard ratios for progression (B) by specific mutation type. Related to Figure 1.

| Mutation Type | Test for larger benefit in tumor volume |  | Conclusion based on 25% False Discovery Rate |
| --- | --- | --- | --- |
|  | p-value | Benjamini-Hochberg critical value for 1-sided test |  |
| L755 Hotspot | 0.031 | 0.025 | N.S. |
| Exon20 Insertion Hotspot | 0.162 | 0.050 | N.S. |
| S310 Hotspot | 0.267 | 0.075 | N.S. |
| V777 Hotspot | 0.276 | 0.100 | N.S. |
| ERBB3 Nonhotspot | 0.421 | 0.125 | N.S. |
| Other Hotspot | 0.529 | 0.150 | N.S. |
| PKD Nonhotspot | 0.651 | 0.175 | N.S. |
| ERBB3 Hotspot | 0.931 | 0.200 | N.S. |
| PKD Hotspot | 0.950 | 0.225 | N.S. |
| Other Nonhotspot | 0.954 | 0.250 | N.S. |

B.
| Mutation Type | Test for larger benefit in hazard ratio for progression |  | Conclusion based on 25% False Discovery Rate |
| --- | --- | --- | --- |
|  | p-value | Benjamini-Hochberg critical value for 1-sided test |  |
| **Exon20 Insertion Hotspot** | **0.011** | **0.025** | **Larger Benefit** |
| S310 Hotspot | 0.094 | 0.050 | N.S. |
| Other Hotspot | 0.274 | 0.075 | N.S. |
| L755 Hotspot | 0.456 | 0.100 | N.S. |
| PKD Hotspot | 0.499 | 0.125 | N.S. |
| PKD Nonhotspot | 0.759 | 0.150 | N.S. |
| V777 Hotspot | 0.944 | 0.175 | N.S. |
| ERBB3 Nonhotspot | 0.962 | 0.200 | N.S. |
| ERBB3 Hotspot | 0.970 | 0.225 | N.S. |
| Other Nonhotspot | 0.985 | 0.250 | N.S. |

**Supplementary Table S4.**
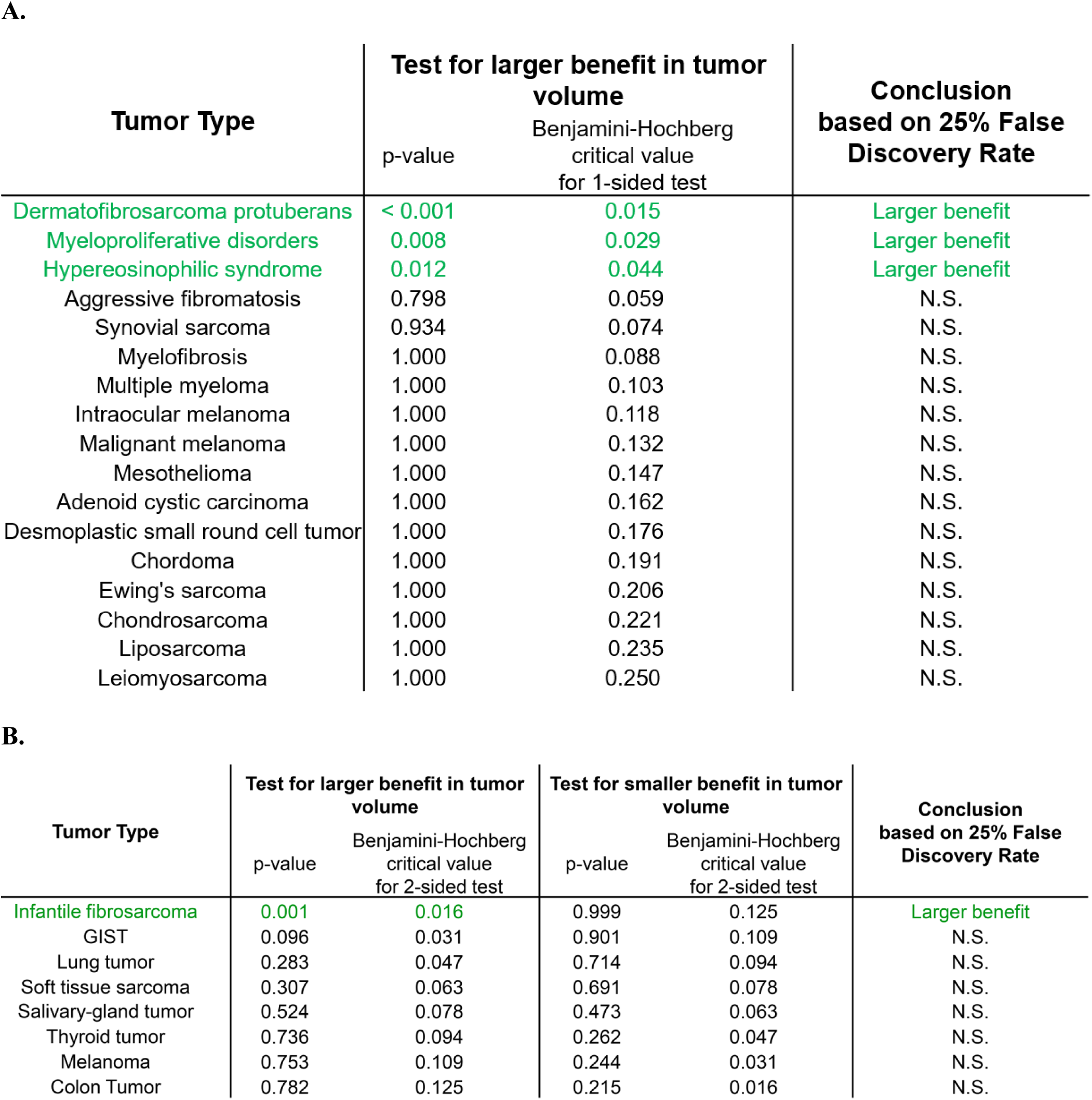
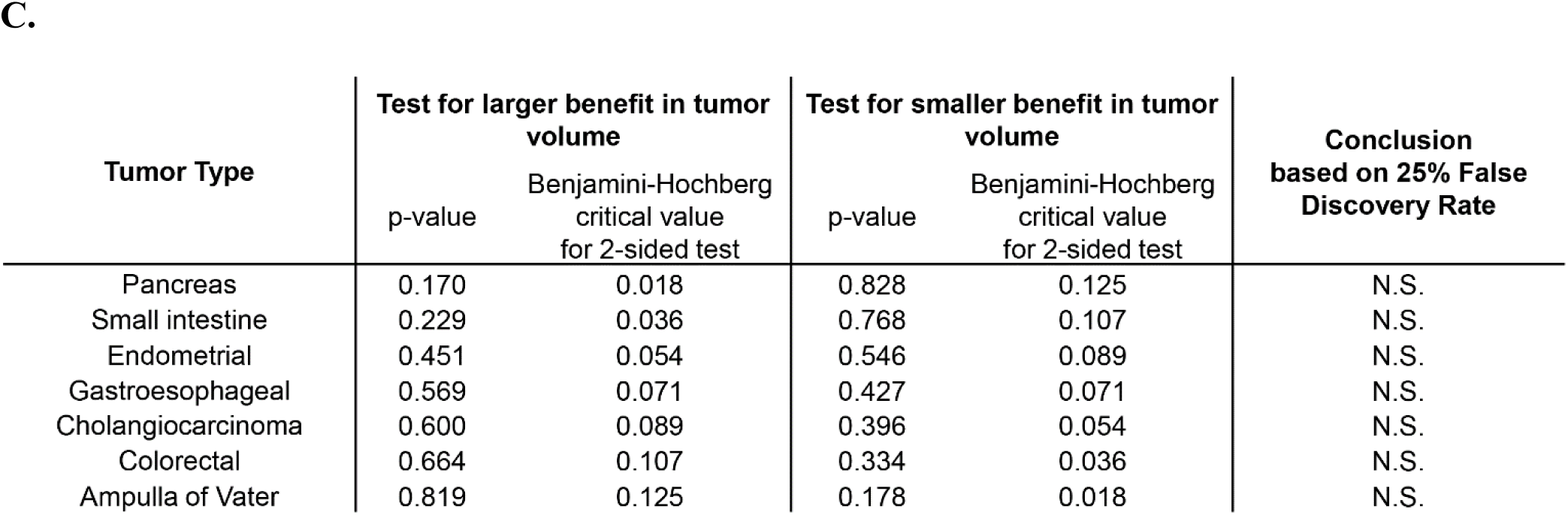
Benjamini-Hochberg critical values for analysis of imatinib (A), larotrectinib (B), and pembrolizumab (C) tumor volume responses by tumor type. Related to Figure 2.

| Tumor Type | Test for larger benefit in tumor volume |  | Conclusion based on 25% False Discovery Rate |
| --- | --- | --- | --- |
|  | p-value | Benjamini-Hochberg critical value for 1-sided test |  |
| Dermatofibrosarcoma protuberans | < 0.001 | 0.015 | Larger benefit |
| Myeloproliferative disorders | 0.008 | 0.029 | Larger benefit |
| Hypereosinophilic syndrome | 0.012 | 0.044 | Larger benefit |
| Aggressive fibromatosis | 0.798 | 0.059 | N.S. |
| Synovial sarcoma | 0.934 | 0.074 | N.S. |
| Myelofibrosis | 1.000 | 0.088 | N.S. |
| Multiple myeloma | 1.000 | 0.103 | N.S. |
| Intraocular melanoma | 1.000 | 0.118 | N.S. |
| Malignant melanoma | 1.000 | 0.132 | N.S. |
| Mesothelioma | 1.000 | 0.147 | N.S. |
| Adenoid cystic carcinoma | 1.000 | 0.162 | N.S. |
| Desmoplastic small round cell tumor | 1.000 | 0.176 | N.S. |
| Chordoma | 1.000 | 0.191 | N.S. |
| Ewing's sarcoma | 1.000 | 0.206 | N.S. |
| Chondrosarcoma | 1.000 | 0.221 | N.S. |
| Liposarcoma | 1.000 | 0.235 | N.S. |
| Leiomyosarcoma | 1.000 | 0.250 | N.S. |

B.
| Tumor Type | Test for larger benefit in tumor volume |  | Test for smaller benefit in tumor volume |  | Conclusion based on 25% False Discovery Rate |
| --- | --- | --- | --- | --- | --- |
|  | p-value | Benjamini-Hochberg critical value for 2-sided test | p-value | Benjamini-Hochberg critical value for 2-sided test |  |
| Infantile fibrosarcoma | 0.001 | 0.016 | 0.999 | 0.125 | Larger benefit |
| GIST | 0.096 | 0.031 | 0.901 | 0.109 | N.S. |
| Lung tumor | 0.283 | 0.047 | 0.714 | 0.094 | N.S. |
| Soft tissue sarcoma | 0.307 | 0.063 | 0.691 | 0.078 | N.S. |
| Salivary-gland tumor | 0.524 | 0.078 | 0.473 | 0.063 | N.S. |
| Thyroid tumor | 0.736 | 0.094 | 0.262 | 0.047 | N.S. |
| Melanoma | 0.753 | 0.109 | 0.244 | 0.031 | N.S. |
| Colon Tumor | 0.782 | 0.125 | 0.215 | 0.016 | N.S. |

| Tumor Type | Test for larger benefit in tumor volume |  | Test for smaller benefit in tumor volume |  | Conclusion based on 25% False Discovery Rate |
| --- | --- | --- | --- | --- | --- |
|  | p-value | Benjamini-Hochberg critical value for 2-sided test | p-value | Benjamini-Hochberg critical value for 2-sided test |  |
| Pancreas | 0.170 | 0.018 | 0.828 | 0.125 | N.S. |
| Small intestine | 0.229 | 0.036 | 0.768 | 0.107 | N.S. |
| Endometrial | 0.451 | 0.054 | 0.546 | 0.089 | N.S. |
| Gastroesophageal | 0.569 | 0.071 | 0.427 | 0.071 | N.S. |
| Cholangiocarcinoma | 0.600 | 0.089 | 0.396 | 0.054 | N.S. |
| Colorectal | 0.664 | 0.107 | 0.334 | 0.036 | N.S. |
| Ampulla of Vater | 0.819 | 0.125 | 0.178 | 0.018 | N.S. |

